# Interneuronal network model of theta-nested fast oscillations predicts differential effects of heterogeneity, gap junctions and short term depression for hyperpolarizing versus shunting inhibition

**DOI:** 10.1101/2022.04.11.487977

**Authors:** Guillem Via, Fernando R. Fernandez, John A. White, Carmen C. Canavier

**Author notes:** Corresponding Author: Carmen C. Canavier.

## Abstract

Theta and gamma oscillations in the hippocampus have been hypothesized to play a role in the encoding and retrieval of memories. Recently, it was shown that an intrinsic fast gamma mechanism in medial entorhinal cortex can be recruited by optogenetic stimulation at theta frequencies, which can persist with fast excitatory synaptic transmission blocked, suggesting a contribution of interneuronal network gamma (ING). We calibrated the passive and active properties of a 100-neuron model network to capture the range of passive properties and frequency/current relationships of experimentally recorded PV+ neurons in the medial entorhinal cortex (mEC). The strength and probabilities of chemical and electrical synapses were also calibrated using paired recordings, as were the kinetics and short-term depression (STD) of the chemical synapses. Gap junctions that contribute a noticeable fraction of the input resistance were required for synchrony with hyperpolarizing inhibition; these networks exhibited theta-nested high frequencies (∼200 Hz) similar to the putative ING observed experimentally in the optogenetically-driven PV-ChR2 mice. With STD included in the model, fast oscillations were only observed before the peak of the theta drive, whereas without STD, they were observed symmetrically before and after the peak. Because hyperpolarizing synapses provide a synchronizing drive that contributes to robustness in the presence of heterogeneity, synchronization decreases as the hyperpolarizing inhibition becomes weaker. In contrast, networks with shunting inhibition required non-physiological levels of gap junctions to synchronize using conduction delays within the measured range; synaptic depression of shunting synapses facilitated fast oscillations, suggesting that shunting inhibition in mEC is desynchronizing.

**Author Summary:** Fast oscillations nested within slower oscillations have been hypothesized to play a role in the encoding and retrieval of memories by chunking information within each fast cycle; networks of parvalbumin positive inhibitory interneurons contribute to the generation of fast oscillations. We show that, in the entorhinal cortex, the intrinsic dynamical properties of these neurons are sufficiently heterogeneous that electrical synapses are likely required to synchronize fast oscillations. Moreover, synchrony likely requires the chemical synapses to have a reversal potential that is negative relative to the action potential threshold of individual neurons during these oscillations. We show that the range of slow phases that support a fast oscillation is controlled by short term synaptic depression. The precise phase locking of the fast oscillation within the slow oscillations is hypothesized to allow for multiplexing of information.

## Introduction

Theta and gamma oscillations in the hippocampus have been hypothesized to play a role in the encoding and retrieval of memories [1–5]. Recent evidence supports the hypothesis that decrements in the temporal precision with which gamma power is coupled to a specific theta phase underlie the decline of associative memory in normal cognitive aging in humans [6]. The medial entorhinal cortex (mEC) generates fast gamma that is thought to convey information about current sensory information to other hippocampal areas [7]. Parvalbumin positive (PV+) neurons are known to contribute to gamma generation; however, the mechanism by which they contribute may differ depending upon the circumstances [8, 9]. In pyramidal interneuronal network gamma (PING) models [10], reciprocal coupling between pyramidal cells and inhibitory interneurons is required to sustain the oscillation, whereas in interneuronal network gamma (ING) models [11], only reciprocal connectivity between inhibitory neurons is sufficient to sustain the oscillation. Recently, it was shown that an intrinsic fast gamma mechanism in mEC can be recruited by optogenetic stimulation at theta frequencies in transgenic mice expressing ChR2 under the Thy1 promoter; Thy1 is expressed in both excitatory and inhibitory neurons [12]. In that study, blocking excitatory transmission abolished theta nested gamma synchrony. However, a more recent study by Butler et al., 2018 [13] in transgenic mice expressing ChR2 under a CaMKIIα promoter found that gamma oscillations were decreased in amplitude but still prominent when excitatory synaptic transmission was blocked. We have also observed fast oscillations nested within optogenetic theta in PV ChR2 mice [14], with presumably little to no contribution from excitatory synapses. Thus, it seems that the contribution of interneuronal interactions to fast oscillations generated in mEC may be variable.

In this study, we have attempted to faithfully capture heterogeneity in the intrinsic and synaptic properties of PV+ fast spiking basket cells in a specific region, the medial entorhinal cortex (mEC). We examine the effect of heterogeneity, gap junctions, synaptic depression, and the synaptic inhibitory reversal potential on synchronization of fast oscillations nested within an excitatory theta drive.

The current study differs from previous studies [11,15,16] on robustness of fast oscillations in inhibitory interneuronal networks to heterogeneity in two principal aspects: 1) the excitability type of the interneuron models and 2) the way in which intrinsic heterogeneity is introduced into the interneuronal network. First, there are two main dynamical mechanisms by which repetitive spiking can arise, corresponding to an early classification of excitability types 1 and 2 [17]. Neurons with type 1 excitability can fire repetitively at arbitrarily slow rates, act as integrators [18], with spiking arising from a saddle node bifurcation [19]. Neurons with type 2 excitability cannot fire repetitively below an abrupt cutoff frequency, act as resonators [18], and their spiking generally arises from a subcritical Hopf bifurcation [19]. Our recent work [20] shows that PV+ fast spiking interneurons in medial entorhinal cortex neurons likely exhibit type 2 excitability, which is consistent with measures also indicating type 2 excitability in striatum [21] and neocortex [22]. Therefore, the model we constructed of the PV+ neurons has type 2 excitability. Second, previous studies used the bias current as the source of heterogeneity but kept the intrinsic dynamics constant. In contrast, in our study, the passive and active parameters of the model were sufficiently variable across the 100 neurons in the network to capture the full range in the experimentally observed membrane time constants and f/I curves. The distinction in the implementation of heterogeneity, along with possible regional differences between mEC and other brain areas, led us to find that, in contrast to the previous modeling results, hyperpolarizing rather than shunting inhibition confers more robustness to heterogeneity.

## Results

### Model Calibration

We first calibrated the passive properties of the model neurons according to experimental data from 11 PV+ cells (see Experimental Methods) as illustrated in Figure 1. Representative voltage traces in response to hyperpolarizing current steps recorded experimentally (Figure 1A) and for a representative model neuron (Figure 1B) are similar., Because the resting membrane potential (RMP) of real neurons ranged from ∼-91 to -69 mV (, Figure 1C1 black boxes), we used values from a similar range for the 100 model interneurons by sampling randomly from a uniform, but narrower, distribution ranging from -81 to -69 mV (red bar) due to the removal of two outliers. The input resistance of the real neurons ranged from ∼ 40-130 MΩ; the values for the model neuron were sampled from a distribution with a slightly larger range of values at larger R_input_. This shift (Figure 1C2) was necessary to reproduce the observed fast firing frequencies, high rheobase currents and steep slopes in the frequency/current (f/I) curves. Experimentally, the membrane time constants τ_m_ ranged from 2.5 to 7.5 ms, with time constants for the model sampled from a uniform distribution between 3 and 7 ms (Figure 1C3). The time constant and input resistance were used to set the membrane capacitance: C_m_=τ_m_/R_input_. These passive properties are consistent with an earlier study [23] .

**Figure 1.**
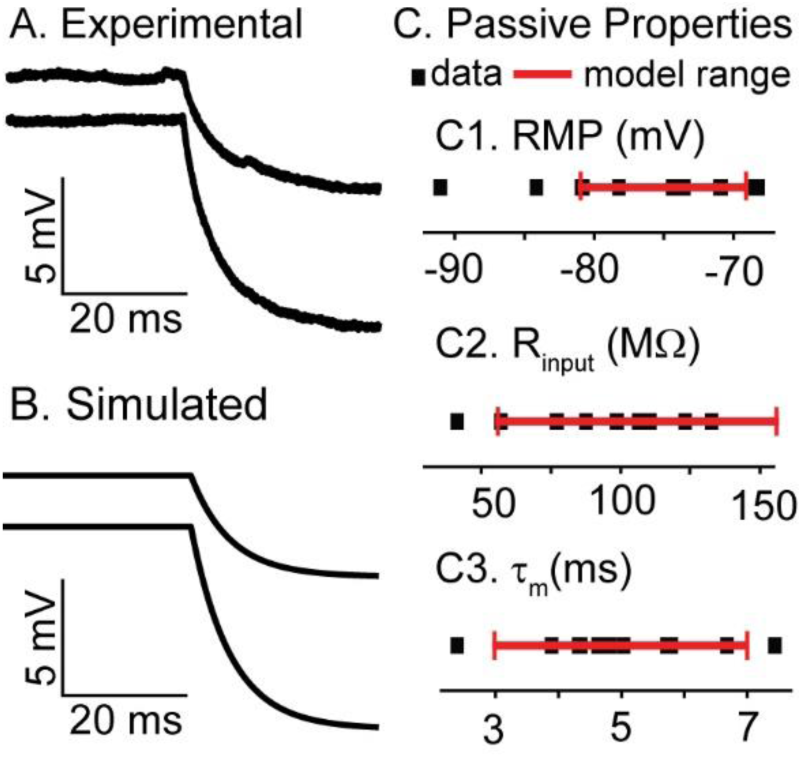
Calibration of passive properties. A. Response of a representative mEC PV+ interneuron to a 50 and a 100 pA hyperpolarizing current step. B. Response of a representative model interneuron to a 50 and a 100 pA hyperpolarizing current step. Parameters were E_L_=-71.79 mV, R_input_=92.07 MOhm and τ_m_=6.4 ms. C. Experimental (black boxes) and model (red bar) ranges of passive properties.C1. Resting membrane potential C2. Input resistance. C3. Membrane time constant.

We next calibrated the parameters for the voltage-gated ionic currents using experimentally recorded voltage traces in response to depolarizing current steps, as described in the Computational Methods. Figure 2A shows representative experimentally recorded voltage traces used to measure the f/I relationship. A step current is applied to determine whether the neuron can support repetitive firing at that level of current. If it can, the stabilized steady frequency is recorded (see Methods) and plotted as the f/I curves in Figure 2C1. The minimum frequencies below which repetitive firing could not be sustained (the cutoff frequency) ranged from 50-200 Hz. Note that below the threshold for sustained, repetitive firing, neurons may fire one or more action potentials before becoming quiescent again.

**Figure 2.**
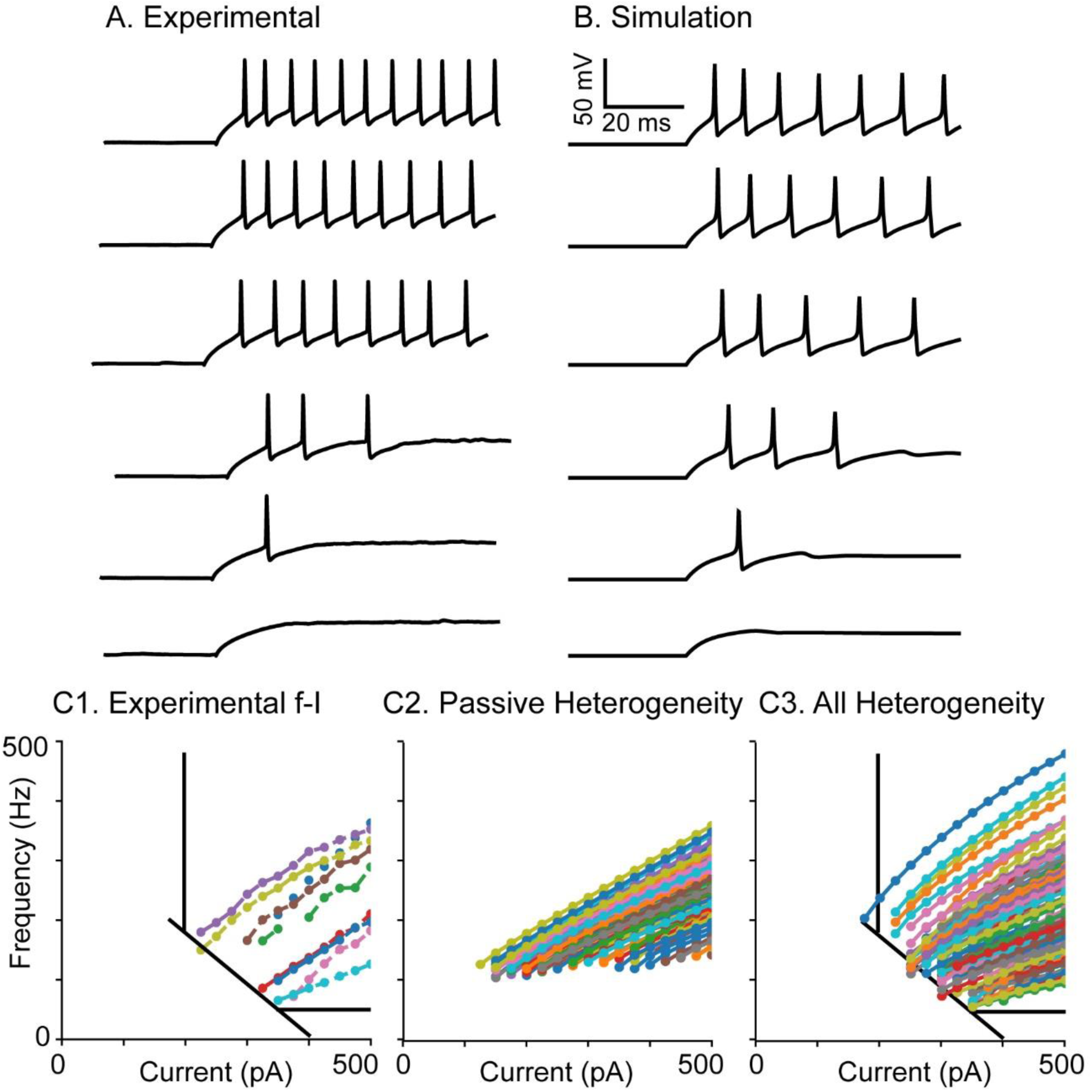
Calibration of active properties. A,B. Responses of representative neurons to depolarizing current steps from 300 pA to 425 pA in 25 pA increments. A. mEC PV+ interneuron . B. Model neuron. Parameters as in Table 1 except E_L_ 78.66 mV, R_input_ 87.86 MΩ (g_L_ 11.38 nS), τ_m_ 5.29 ms (C_m_ 0.060 nF), g_NA_ 18930 nS, g_Kv1_ 48.79 nS, g_Kv3_ 784.56 nS, θ_m_ -45.78 mV, θ_h_ -53.62 mV, θ_n_ 21.97 mV, θ_a_ = 46.55 mV. C. Population f-I curves. C1. Experimental. C2. Passive heterogeneity only, with active parameters fixed at, g_NA_ 16522.9 nS, g_Kv1_ 76.3 nS, g_Kv3_ 705.5 nS, θ_m_ -56.42 mV, θ_h_ -56.59 mV, θ_n_ -4.23 mV, θ_a_ = 51.4 mV. C3. Heterogeneity in both active and passive properties.

**Table 1.** Parameters for gating variables. The θ parameters are given for the homogeneous network in Figure 4. These parameters were varied across the network in order to reproduce the variability in f/I curves. The other parameters were held constant for all model neurons.

| | $m$ | $h$ | $n$ | $a$ |
| --- | --- | --- | --- | --- |
| $\theta$ | -56.42 | -56.59 | -4.23 | 51.40 |
| $\sigma_1$ (mV) | 4 | -20 | 12 | 12 |
| $\sigma_2$ (mV) | -13 | 3.5 | -8.5 | -80 |
| $k_1$ (ms<sup>-1</sup>) | 0.25 | 0.012 | 1 | 1 |
| $k_2$ (ms<sup>-1</sup>) | 0.1 | 0.2 | 0.001 | 0.02 |

The parameters for the Na and Kv3 currents were calibrated (see Methods) in the absence of Kv1 to reproduce the action potential waveform. Inclusion of Kv1 in the model was necessary to allow firing one or more spikes before becoming quiescent, as well as reproducing the variability in the neuron cutoff frequencies. This is consistent with previous studies showing that Kv1 and Kv3 set the minimum and maximum firing rates, respectively. [24, 25]. A representative model neuron in Figure 2B reproduces the transient responses after current step onset at near-rheobase current amplitudes (compare to Figure 2A). The action potentials have features similar to the recorded neurons, including spike height, width, and after-hyperpolarization depth. Figure 2C2 shows the f/I curves for model neurons with heterogeneous passive properties but homogeneous active parameters, corresponding to a neuron near the center of the spread in Figure 2C1. The cutoff frequencies were far too uniform when only passive properties were varied. Therefore, the peak conductances and the half-activation and inactivation voltages for the voltage-gated currents were made heterogeneous across the 100 neurons (see Methods). We eliminated any parameter sets that resulted in an f/I curve too far outside the envelope indicated by the black lines in Figure 2B1 and B3, or that did not preserve other desired features (see Methods). The model network with heterogeneity in both active and passive properties was able to reproduce the broad distribution of f/I curves, as shown in Figure 2C3. Overall, the qualitative agreement between model and data is good both in terms of action potential shape and steady-state firing frequencies. The intrinsic parameters were frozen at the values that generated Figure 2C3, and only connectivity parameters were varied in the subsequent simulations. In Figure 3, the experimental histograms, and fits for the distribution of peak conductances in chemical and electrical synapses are shown, with parameters and connection probabilities given in the Methods section.

**Figure 3.**
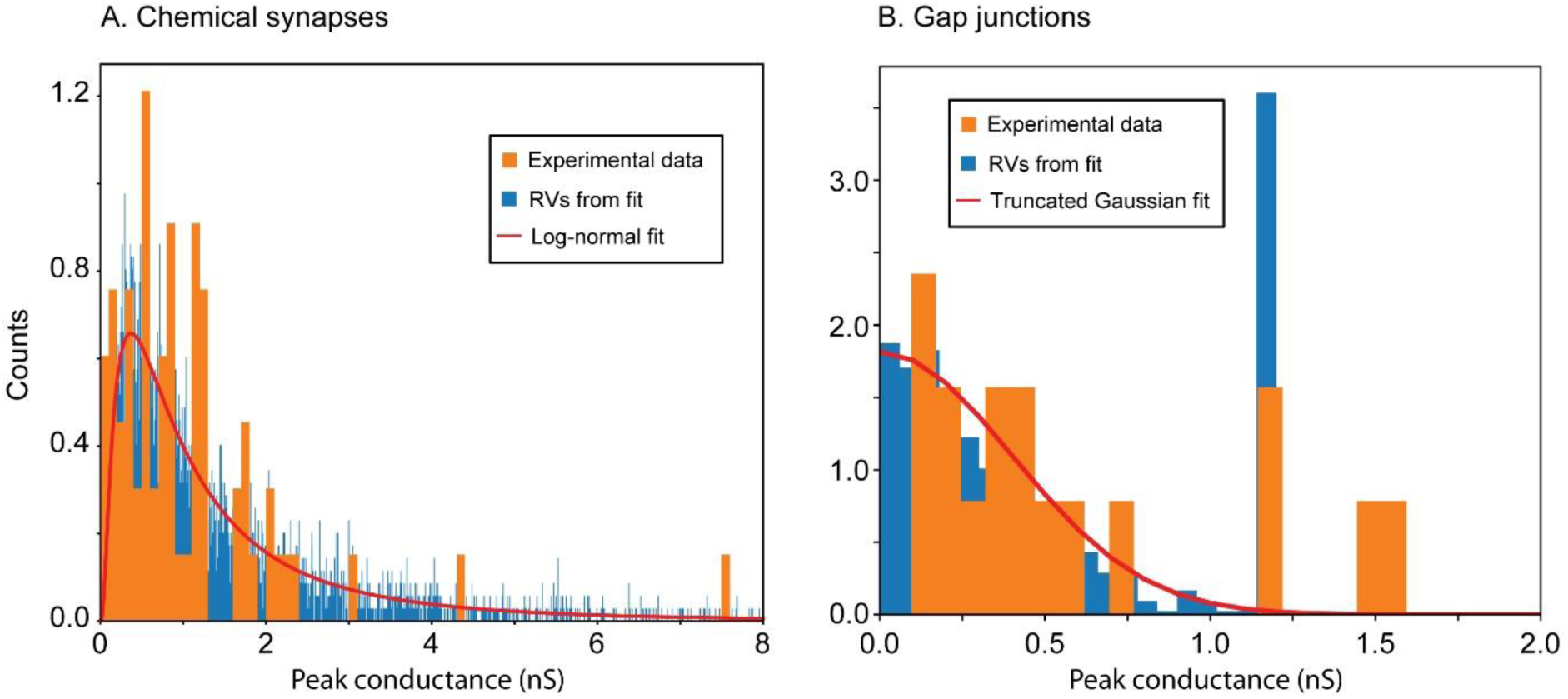
Calibration of synaptic properties. A. Chemical Synapses. B. Electrical Synapses.

### Response of Homogeneous Networks with Hyperpolarizing versus Shunting Synapses to Theta Modulation: Phase Response Curve Analysis

After calibrating the model network, we assessed its response to a theta-modulated external input that simulates an optogenetic protocol to study its synchronizing properties. To gain theoretical insights into the model, we first considered homogeneous networks amenable to analysis via phase response theory under the assumption of pulsatile coupling [26–28]. The homogeneous network consisted of 100 clones of a single model neuron, with identical intrinsic properties, connected through 36 identical, but randomly assigned, presynaptic chemical synapses. The parameters of the single model neuron correspond to one with an f-I curve close to the middle of the heterogeneous range. The active parameters for voltage-gated ion currents were the same as in Figure 2C2, and the passive parameters were set to τ_m_=4.52 ms, E_L_=-70.38 mV and g_L_=15.35 nS. We incorporated synaptic depression at the chemical synapses, but we did not include gap junctions in order to determine whether the chemical synapses alone could synchronize the network.

Chemical synapses are modeled as GABA_A_ synapses. Their reversal potential is difficult to measure *in vivo*; it is unclear whether they are hyperpolarizing or shunting (see Discussion). In order to generate testable predictions that differ depending on whether the synapses are shunting or hyperpolarizing, we compared model network dynamics using synaptic reversal potentials, E_syn_, values of -75 mV (left column in Figure 4) and -55 mV for hyperpolarizing and shunting synaptic inhibition, respectively (right column in Figure 4). Although shunting networks exhibit faster and more variable frequencies in response to simulated optogenetic sinusoidal drive, both exhibit global synchrony. Since the neurons are identical and receive identical input, no connectivity is actually required to synchronize them. However, the perfect synchrony present with chemical synapses intact indicates that the synapses themselves are not sufficient to destabilize global synchrony in the presence of a common sinusoidal drive. Adding the full heterogeneity to the intrinsic properties of the neurons in the network as described in Figures 1 and 2 completely eliminates the fast oscillations nested in the theta drive (Figures 4A2, B2). In Figures 4A3 and B3, we show the response for a network using identical neurons, with heterogeneity only in the synaptic delays, weights, and numbers of presynaptic partners. For hyperpolarizing inhibition, the height of the first peak is 100, indicating global synchrony for all 100 neurons as in Figure 4A1. However, the peaks widen and diminish in height until fast oscillations disappear. The synchronization properties of the same network using shunting chemical synapses also decays but not as abruptly.

**Figure 4.**
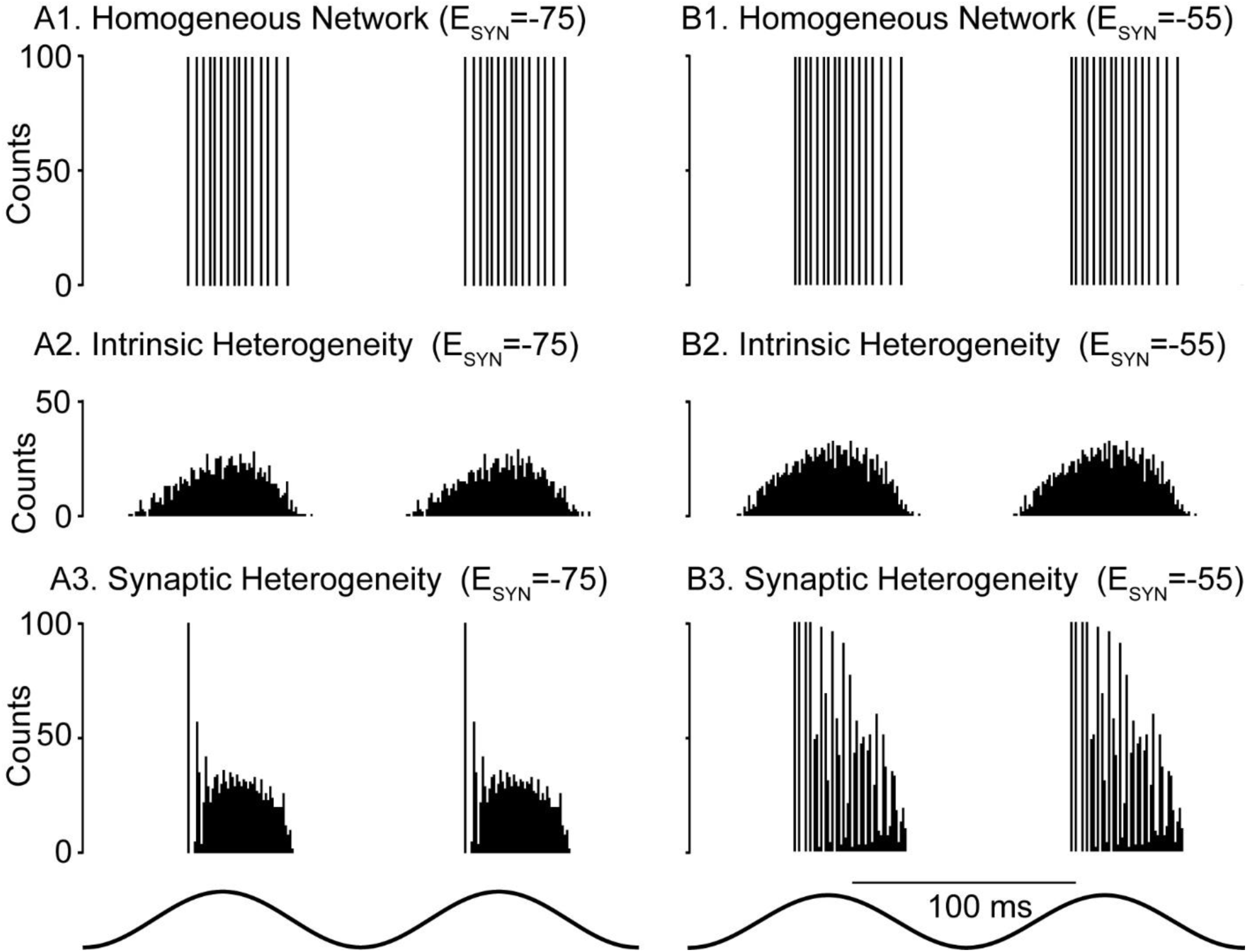
Biophysically calibrated levels of heterogeneity are desynchronizing. Representative spike histograms as a function of time. A1 Hyperpolarizing and B1 Shunting Homogeneous Networks with these parameters: g_NA_ 16500 nS, g_Kv1_ 76.3 nS, g_Kv3_ 705.5 nS. E_L_ -70.4 mV and C_M_ 0.069 nF. g_L_ was 15.3468 nS resulting in an input resistance of 65.16 MΩ. Each neuron received exactly 36 chemical synapses with a strength of 1.65 nS. The synaptic delay was fixed at 0.8 ms. The optogenetic theta drive (bottom) varies sinusoidally at 8 Hz from 0 to 14 nS.

Next, we used a mean field approach that assumes synchronized oscillations in a homogeneous network in which every neuron is identical and receives identical input (i.e. from exactly 36 other identical neurons). This allowed us to perform a phase response curve (PRC) analysis to predict the stability and frequency of global synchrony [26–28]. In order to apply phase response theory, an autonomous system with constant parameters is required. Therefore, we considered the case of a constant external input at the midpoint of the theta modulation of g_ChR_ (7 nS) (Figure 4). Under these conditions, the intrinsic frequency of the model neuron is 210 Hz. We generated a phase response curve by applying an inhibitory postsynaptic conductance on separate trials at each of 100 equally spaced phases within the free-running cycle of the model neuron, with the point at which the neuron reaches threshold (defined as -30 mV) taken as a phase of 0 (and 1). We used a conductance that was 36 times larger than the conductance of a single synapse to simulate the synchronous input received by a single neuron in the network during global synchrony (see inset in Figure 5A). We plotted the normalized increase in the period (the phase delay) as a function of the phase (Figure 5A) for a hyperpolarized synaptic reversal potential (red trace) and for a shunting one (solid green trace). Hyperpolarizing inhibition (red) consistently induces phase delays that lengthen the period, whereas shunting inhibition (green) consistently advances the phase (negative phase resetting in our convention) and shortens the period. Thus, we expect higher frequencies for networks with shunting rather than hyperpolarizing synapses. The two dotted lines indicate the range across which delays were varied in Figure 4C. The leftmost arrows indicate the phase at which an input would be received in a globally synchronous mode at: θ=δ/P_i_, where P_i_ is the free running period of the neuron. For a delay of 0.8 ms, this phase is 0.168.

**Figure 5.**
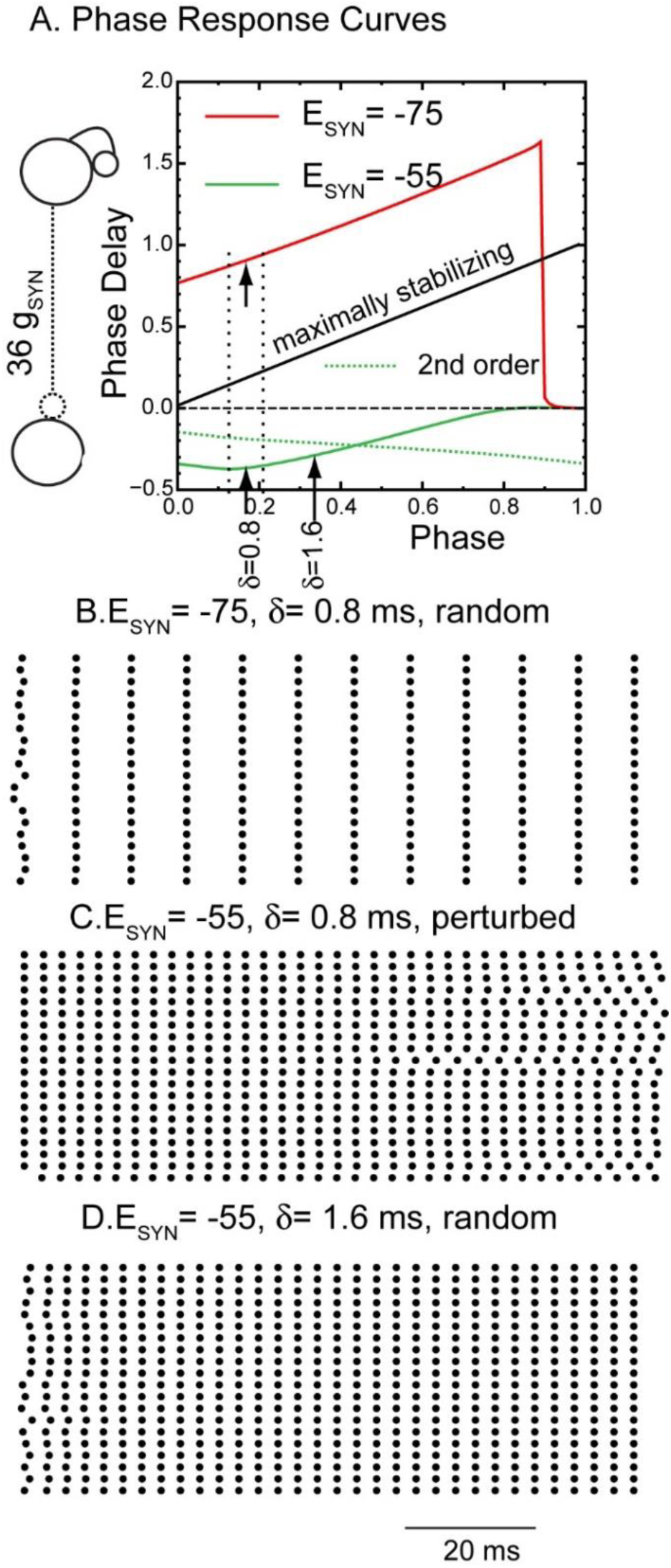
Phase Response Curves Explain Synchronizing Tendencies. A. A biexponential inhibitory postsynaptic conductance as the perturbation to a single neuron from Fig. 4A to generate the PRC for hyperpolarizing (red) and shunting inhibition (green). The dashed green curve shows the normalized change in the cycle after the cycle that contains the perturbation (second order). The strength of the individual conductance was multiplied by 36 to reflect the 36 simultaneous inputs received by a single neuron (left inset) during perfectly synchronous oscillations. The leftmost arrows indicate the phase at which an input delayed by 0.8 ms is received in the network. The dashed lines refer to the range of synaptic delays shown in Figure 4C. The free running period of this neuron is 4.7625 ms at a constant ChR conductance of 7 nS, the midpoint of the excitatory theta drive. B. For hyperpolarizing synapses with conduction delays of 0.8 ms, synchrony is stable and attracts from random initial conditions in a single cycle. C. For shunting synapses, starting from exact synchrony, perturbing even a single neuron (bottom trace) eventually desynchronizes the network. D. If the conduction delay is increased to 1.6 ms in the network with shunting inhibition, synchrony is stabilized and attracts quickly from random initial conditions. B-D are raster plots of 20 representative neurons from the 100 neuron network.

In order to apply the phase response curves, an inhibitory input must have the same effect in the network as it did when applied to a free-running neuron at a stabilized frequency in order to generate the PRC in the first place. In practice, the requirement is that the neuron must have returned very close to its unperturbed state (on its limit cycle, speaking mathematically) by the time the next input is received. If the second order phase response [29, 30], that is, the change in length in the subsequent cycle, is small, then it is likely that this requirement is satisfied.For delays up to 90% of the cycle period, there is less than a 3% change in the length of the subsequent cycle (not shown). The phase resetting resulting from an input applied at a delay of 0.8 ms predicts a network period of 110 Hz for hyperpolarizing synapses. Global synchrony in the homogeneous network is strongly attracting; the network converges to global synchrony in a single cycle after random initialization (Figure 5B). The observed frequency in the homogeneous network is 115 Hz, which is not exact, but is very close to the predicted frequency and illustrates the predictive power of the theory despite some slight deviation from the pulsatile coupling assumption.

Phase response theory can also explain the fast convergence to synchrony. The stability of synchrony is determined by whether a perturbation of even a single neuron from the globally synchronous mode decays or grows on the next cycle. For a synchronous mode with short delays, the perturbation grows or decays according to the scaling factor 1−*f*_3_′_6_ (*θ*) − *f* ′(*θ*) [26, 28], where *f*_3_′_6_ (*θ*) is the slope of the phase response curve of the single perturbed neuron at the locking phase (arrow on green curve in Figure 5A) and *f* ′(*θ*) is the slope of the phase response curve of the other 99 neurons in response to an input from the perturbed single neuron, also at the locking phase. The other 99 neurons are assumed to receive 35 simultaneous inputs with a delay of 0.8 ms after the population spike, with the perturbed neuron only adding a 36th simultaneous input to the already very strong input, allowing us to neglect *f* ′(*θ*) as small compared to *f*_3_′_6_ (*θ*) . The expression for the rate at which a perturbation decays then becomes approximately 1− *f*_3_′_6_ (*θ*). The maximally stabilizing value is *f*_3_′_6_ (*θ*) = 1 (see illustrative maximally stabilizing diagonal in Figure 5A) since after only a single cycle a perturbation will decay to zero. The slope of the PRC for hyperpolarizing synapses (arrow on red trace) is close to one, which accounts for the rapid convergence in Figure 5B. The sharp decrease in the PRC at late phase occurs because of the finite duration of the waveform of the biexponential synaptic conductance; when it is applied near the end of the cycle, insufficient charge accumulates to delay the action potential substantially; instead, it lengthens the subsequent cycle length. However, if an input is initiated after the action potential has occurred, the cycle containing the start of the perturbation is substantially lengthened. The discontinuities near a phase of 0.9 for the red trace and between 1 and 0 for both the red and solid green trace in Figure 5A are highly destabilizing. For global synchrony to be observed in networks with any jitter in the spike times, the conduction delays must be long enough (and short enough) to avoid sampling a discontinuity [31].

The PRC approach is less informative for the network with shunting inhibition because the pulsatile coupling assumption is not well honored. Shunting inhibition decreases the network period, such that the synaptic waveform due to single input persists throughout two or more cycles. Therefore, we plotted the change in cycle length in both the cycle that contains the start of the perturbation (first order PRC, solid green curve in Figure 5A) and in the second cycle (second order PRC, dotted green curve). The slope of the first order PRC is initially negative, which means that very short delays <0.6 ms are destabilized because a negative *f*_3_′_6_ (*θ*) causes the perturbation multiplier1− *f*_3_′_6_ (*θ*) to have an absolute value greater than one, which leads to growth of the perturbation. Flat PRCs with a slope near zero are only very weakly attracting or repelling. We initialized all neurons but one in the shunting network on their unperturbed, free-running steady limit cycle at the action potential threshold in Figure 5C. The remaining neurons were initialized at a phase that corresponded to a difference of one tenth of the period relative to how the other neurons were initialized. This slight perturbation slowly desynchronized the network, which demonstrates that shunting inhibition at a delay of 0.8 ms (left arrow) is destabilizing for global synchrony. The theta-modulated synchrony in Figure 4B for the shunting network likely occurs in spite of, and not because of, the weakly desynchronizing chemical synapses and is therefore driven solely by the common input to identical neurons. Although phase response theory under pulsatile coupling does not strictly apply to the shunting networks, it can still provide some insights. For example, it suggests that the network will synchronize if the conduction delay is increased from 0.8 ms to 1.6 ms (rightmost arrow on solid green curve), where the slope is closer to one and more strongly synchronizing. Indeed, synchrony arises from random initial conditions (Figure 5D) when the conduction delay is set to 1.6 ms. The convergence, however, is not as fast, which is likely due to the slope being farther from one. The observed frequency of 325 Hz is again similar to the predicted frequency of 295 Hz. The first order PRC for shunting, but not hyperpolarizing, inhibition reverses sign at a phase of 0.13, very near the action potential trough at a phase of 0.15. This suggests that action potential width, along with conduction delay and synaptic rise time, may be a determinant of synchronization tendencies for shunting, but not hyperpolarizing inhibition in the mEC. The shunting PRC is relatively flat between the two dotted vertical lines that indicate the range of variability in the conduction delays in Figure 4C. The relative insensitivity to variability in the conduction delay implies greater robustness of the shunting network to variability in synaptic delays compared to the hyperpolarizing network, which likely contributes to the greater robustness of shunting networks to variability in both synaptic delays and synaptic conductances shown in that figure.

### Response of Heterogeneous Networks with Hyperpolarizing versus Shunting Synapses to Theta Modulation: Effects of Gap Junctions and Synaptic Depression

We showed in Figure 4B that the full complement of observed heterogeneity in the intrinsic properties of the model by itself suppressed theta-nested fast oscillations. Therefore, it is unsurprising that in the presence of full heterogeneity (both intrinsic and synaptic), theta-nested fast oscillations are also suppressed in networks with either hyperpolarizing or shunting inhibition (Figure 6A1 and B1). However, the heterogeneous networks in Figure 4 B and C neglected gap junctions in order to assess the effects of chemical synapses in isolation. In fact, gap junctions between PV+ interneurons in the mEC are highly prevalent [32], therefore we tested their impact in the model. As described in the Methods, the experimentally recorded connection probability and distribution of gap junction peak conductances suggest that they make a substantial contribution to the measured input resistance. Our initial calibration of f-I curves in Figure 2C ignored gap junctions; therefore, incorporating gap junctions required recalibration of the passive properties of the model neurons. As described in the Methods, we reduced the leakage conductance and adjusted the reversal potential each time a gap junctional conductance was added to a model neuron in order to preserve the original range of values for the input resistances and resting membrane potentials. A minimum value for the leak conductance, *g_L_^min^*, was set at 1.5 nS to honor the constraint that the interneurons must have at least some intrinsic leak conductance. Although imposing this constraint decreased the total number of electrical synapses, the effect was mitigated because the exact number of synaptic contacts of either kind is not known in the mEC, as noted in the Methods. The parameters for voltage-gated ion currents were not changed, and f/I curves were minimally affected by adding the gap junctions (Figure S1).

**Figure 6.**
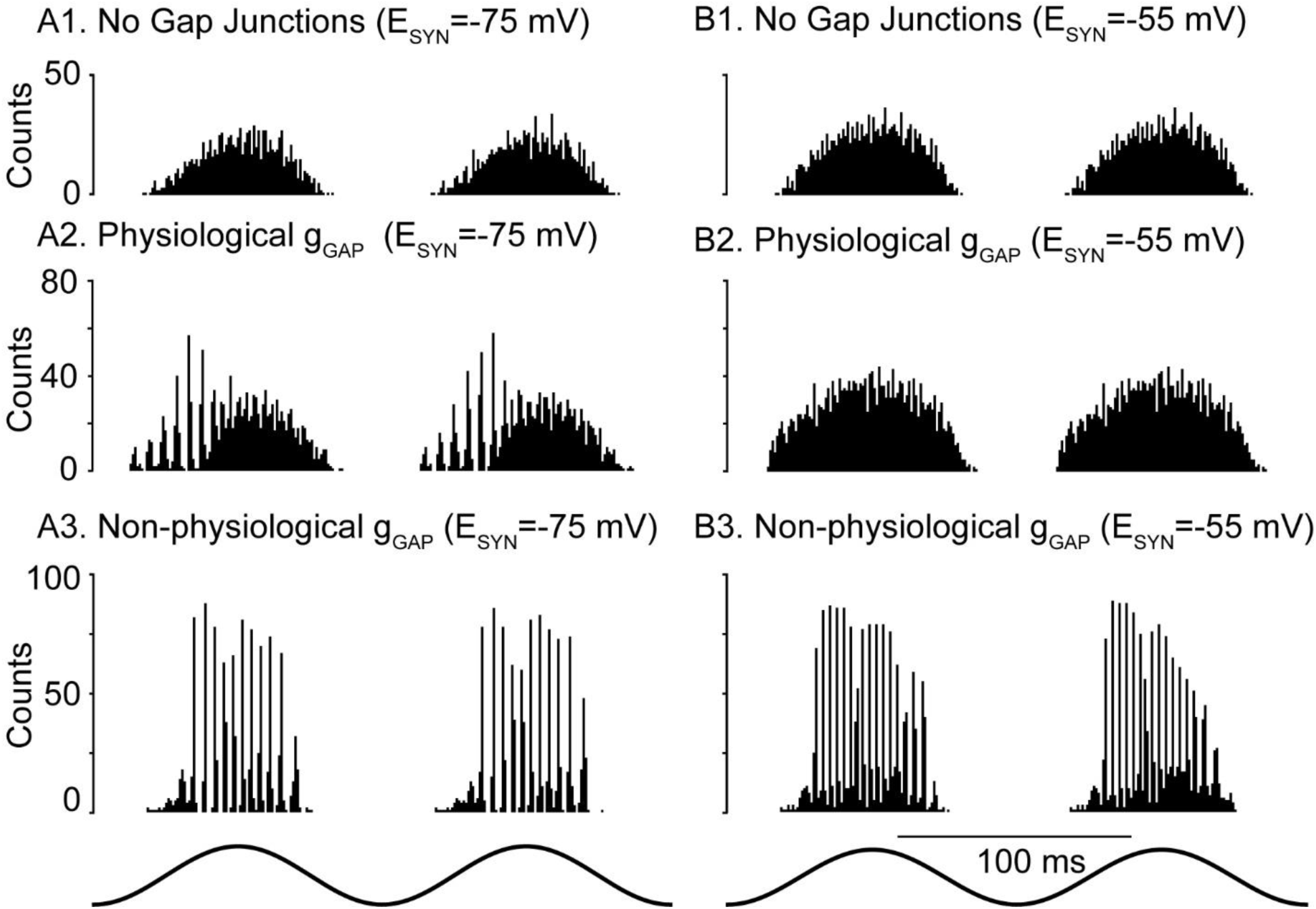
Gap junction connectivity is required for theta nested fast oscillations in heterogeneous networks. Representative spike histograms as a function of time. A. A Fully heterogeneous networks (both intrinsic and synaptic heterogeneity) with no gap junctions. A1. Hyperpolarizing. A2. Shunting. B. Heterogeneous networks with gap junctions calibrated according to Figure 3 and he text accompanying this figure. B1. Hyperpolarizing. B2. Shunting. C. Heterogeneous networks with 2 nS gap junctions with no compensatory reduction in leakage conductance. C1. Hyperpolarizing. C2. Shunting.

For networks with hyperpolarizing chemical synapses, when physiologically constrained gap junctions were added as described above, network synchrony at fast frequencies built up slowly during the theta cycle (Figure 6A2), then decreased as in the homogeneous network (Figure 4C1). Fast oscillations were not visible for physiological gap junction connectivity with shunting chemical synapses (Figure 6B2). Strong homogeneous gap junction strengths (2 nS) resulted in tight synchronization for both hyperpolarizing and shunting inhibition (Figures 5A3 and B3). Under these conditions, however, it was not possible to compensate for such strong gap junctional conductances by decreasing the leak conductance while still maintaining input resistance values within the experimentally constrained ranges. Also, this strength is beyond the physiologically observed range in Figure 3. We deemed this gap junctional connectivity non-physiological for those two reasons. The greater robustness of global synchrony of theta-nested fast oscillations was preserved at the larger value (1.6 ms) of conduction delay (Figure S2), despite the synchronizing tendency of networks with shunting synapses detected at that value in Figure 5C.

In order to demonstrate that the results in Figure 6 were not specific to one random connectivity pattern, we constructed 30 networks that differed in their connectivity pattern in both chemical and electrical synapses. Besides the distinct connectivity graphs, the peak conductances for the two types of synapse and the delays for the chemical ones were obtained from a different sampling of their respective distributions. In Figure 7, each filled dot represents a different network, and the physiological level of gap junctional connectivity was implemented as in Figure 6A2. Physiological levels of gap junctions were effective in stabilizing networks using hyperpolarizing synapses (Figure 7) but not shunting synapses, regardless of the connectivity pattern. No evidence for nested fast oscillations was found for shunting networks; hence, there is no corresponding plot for that case. The frequency of the nested oscillations is relatively stable across network instantiations for hyperpolarizing synapses (Figure 7). The results in Figure 7 were qualitatively preserved when a truncated Gaussian fit to the chemical synaptic conductance strengths was used instead of the lognormal distribution (not shown). The different degrees of synchrony for different connectivity patterns suggest that higher order statistics in the connectivity graph, like the presence of hubs or loops, could also enhance or hinder synchrony. Alternatively, networks in which similar neurons are more strongly connected might be more predisposed to synchrony.

**Figure 7.**
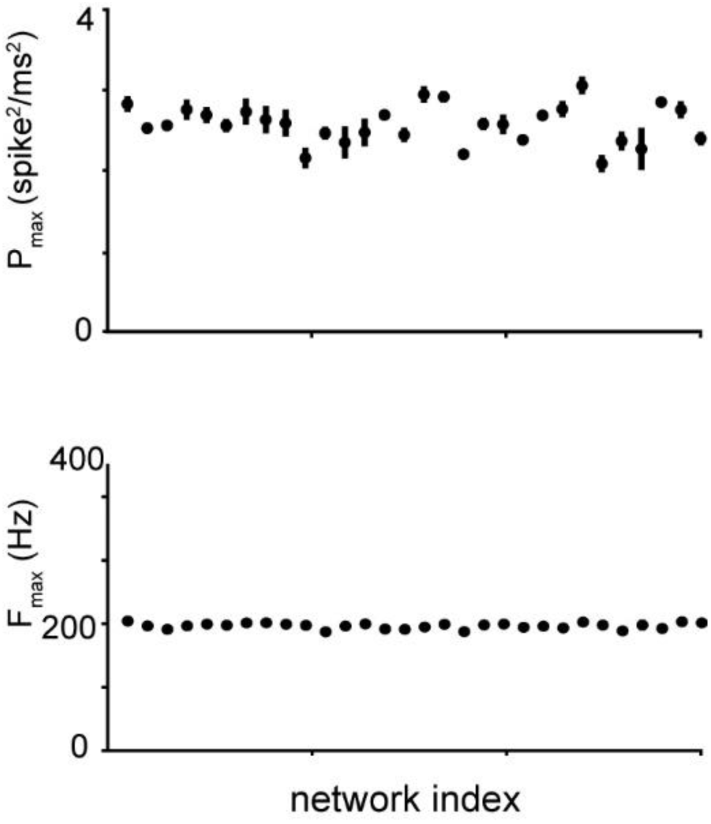
Performance of hyperpolarizing network is robust to different instantiations of network connectivity. The fast frequency with the most power is shown at the bottom for each of 30 networks, and the power at that frequency is given at the top. There is some variability across theta cycles within a given network, indicated by the vertical lines showing the standard deviation.

We next examined the effect of short-term depression (STD) on the model networks by removing STD from the model. For networks with hyperpolarizing synapses, we previously showed (Figure 6A2) that they preferentially exhibited fast oscillations on the intracellular rising (extracellular falling) phase of theta stimulation compared to the falling phase. Removing STD clearly negated this phase preference (compare Figure 8A2 to 8A1) and rendered the fast oscillation amplitude symmetric about the peak (extracellular trough). Since we were unable to obtain theta-nested fast oscillations in the shunting networks without unrealistically strong gap junctions, we tested the effect of removing STD on highly synchronous networks with excessive gap junctional coupling, which we previously showed produced synchrony (Figure 6B3). Removing STD did not change the phase preference, but clearly decreased the amplitude of the fast oscillations. Since STD decreases the overall contribution of the chemical synapses, we conclude that in our networks, hyperpolarizing inhibition increases the robustness of synchrony at fast frequencies, whereas shunting inhibition decreases it. In fact, increasing the strength of the shunting conductance greatly decreased the power of the fast oscillations with very strong gap junctions (Figure S3). Figure 8C1 shows a scalogram for a representative network with STD showing that the power is concentrated in the 150-200 Hz range. The onset and offset of fast oscillations were computed from the phases at which the power crossed from below and from above, respectively, a threshold of 0.3 times the maximum power across the 30 cycles of the simulation. The circular histograms in Figure 8C2 show these onset (blue) and offset (red) phases pooled across 30 simulated theta cycles of all 30 simulated networks with different connectivity patterns. The case with STD is shown on the left, and without STD on the right. The offset phase (in radians) with and without STD was -1.171±0.002 and 1.649±0.003, respectively, with zero being the peak of the theta stimulation. Further, the onset phase with and without STD was -2.406±0.002 and - 2.268±0.002, respectively. The theta phase offset was substantially and significantly different (p<0.001) using Watson’s U2 test in the circular statistics package in R (CRAN, RRID:SCR_003005) [33]. The theta phase onset differed only very slightly between the two conditions; however, this difference was also significant (p<0.001). The smaller range of theta phases that support nested fast oscillations with STD compared to without them supports the premise that hyperpolarizing inhibition helps to synchronize inhibitory networks in the presence of biological levels of heterogeneity, provided there is also a biological level of gap junctional connectivity.

**Figure 8.**
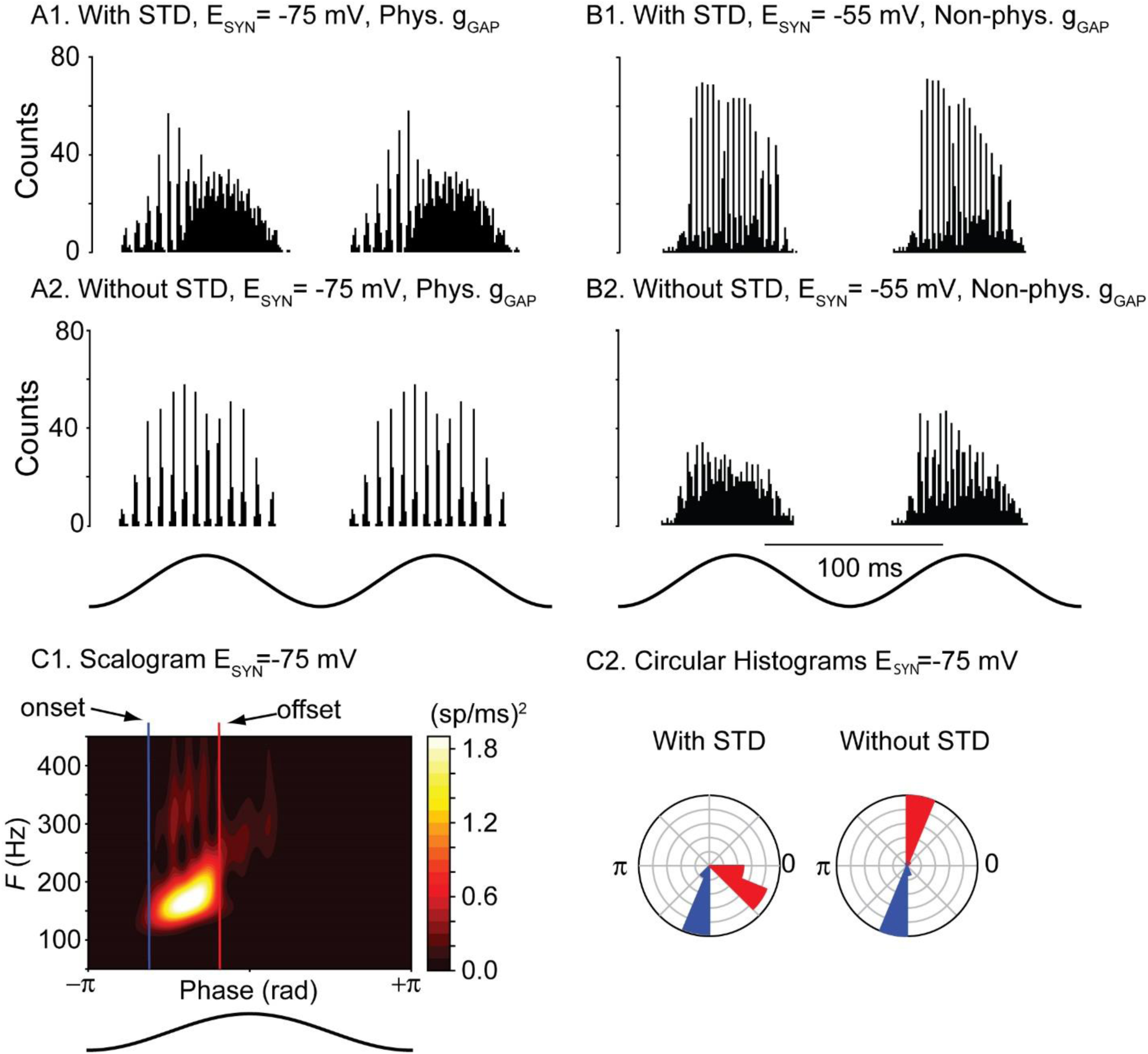
STD induces a preference for theta phases before the peak in hyperpolarizing but not shunting networks. A1. Repeated from Fig. 7A2, theta-nested fast oscillations increase on the rising phase of theta stimulation but decrease after the peak due to synaptic depression. B1. Repeated from Figure 6B3. Theta-nested fast oscillations are more symmetric with respect to the peak. A2. Removing short-term depression from the network restores symmetry about the peak for hyperpolarizing networks. B2. Removing short-term depression from the network decreases synchrony but does not appear to affect symmetry about the peak for shunting networks. C1. Scalogram showing how onset and offset phases were determined. C2. Circular histogram of onset (blue) and offset (red) phases with and without STD.

## Discussion

### Summary

We used an experimentally calibrated computational model of a network of fast-spiking parvalbumin-positive inhibitory basket cells (PVBCs) to study their synchronizing properties, as well as the properties of the emerging oscillations and the underlying mechanisms. The model was calibrated using electrophysiological recordings from mouse mEC slices and reproduced the full range of heterogeneity in the experimental f/I curves, including their high cutoff frequencies, which indicate type II excitability. We calibrated both the neural intrinsic passive and active properties. The former includes the leakage reversal potential, leakage conductance and membrane time constant. The latter comprise the parameters that determine the dynamics of voltage-gated ion currents. We also used recordings to calibrate the properties of both chemical and electrical synapses between the neurons. Network behavior was studied using a theta-modulated input current simulating a channelrhodopsin-driven optogenetic input similar to previous studies [12–14].

Our results suggest that a preference for fast oscillations in the rising phase of excitatory theta drive (descending phase of extracellular theta) is a hallmark of hyperpolarizing inhibition when combined with the short-term synaptic depression commonly observed in synapses made by PV+ neurons [34, 35]. This phase preference is consistent with that observed in PV+ basket cells in CA1 [36]. Based on STD parameters in PV+ mEC interneurons [32], this preference will be expressed even at the lower range of fast gamma in mEC (∼65 Hz). Our results strongly suggest that any ING in mEC is mediated by hyperpolarizing rather than shunting inhibition, and that artificially manipulating the synaptic reversal potential to make it more depolarizing should decrease optogenetically-evoked ING that persists after blocking excitatory synapses. As in a previous study [37], we observed synergy between gap junctions and hyperpolarizing inhibition. Gap junctions mitigate the effect of heterogeneity by forcing the activity of different neurons to be more similar than if gap junctions are absent. Further predictions are that gap junctional connectivity is required for the expression of fast oscillations mediated by inhibitory interneurons in the mEC, and that blocking gap junctions would not only disrupt synchrony, but also change the measurable input resistance and time constants of isolated PV+ neurons. This is consistent with previous estimates [38] in which gap junctions account for one third to one half of the input conductance of fast spiking interneurons. Unfortunately, blocking gap junctions selectively is challenging experimentally as the gap junction blockers are non-specific and can block voltage-gated K^+^ currents, which limits the ability to discern the specific impact of gap junctions on input resistance [39].

### Shunting versus Hyperpolarizing Synapses

Here, we modeled chemical synapses between PV+ cells as ionotropic GABA_A_ receptors. Whether inhibition is shunting or hyperpolarizing depends upon the chloride reversal potential, as well as on the reversal potential for bicarbonate ions flowing in the opposite direction [40, 41]. Larger contributions of bicarbonate lead to more depolarized synaptic reversal potentials, and these contributions may vary between brain regions. Early studies in CA1 and CA3 found hyperpolarizing inhibition between basket cells [17, 18]; in contrast, the inhibition between FS basket cells in the dentate gyrus *in vitro* is shunting, with a reversal potential of about -52 mV [15]. Moreover, the intracellular Clconcentration is not static and can be modulated; for example, activation of kainate-type glutamate receptors potentiates the activity of the potassium-chloride co-transporter 2 (KCC2) via interactions with the GluK2 subunit, reducing the intracellular chloride concentration and rendering the reversal potential more hyperpolarized [42]. There is also evidence that steady-state intracellular Cl– gradients within neurons may be set by cell-type-specific, subcellular expression patterns of functional cation chloride cotransporters [41]. As a result of all these factors, there is sufficient uncertainty regarding the precise reversal potential of GABA_A_ synapses between interneurons to warrant our systematic study of both types of inhibition.

### Fast Oscillations in the Entorhinal Cortex

At least two types of fast oscillations have been observed in the entorhinal cortex: fast gamma oscillations in the 65-140 Hz range [7] and ripples 140-200 Hz [43]. The frequency border between these oscillations is ambiguous in the literature; some quote 100 Hz as the border [44], whereas others label 90-150 Hz as an epsilon band or alternatively refer to the 65-90 Hz range as medium gamma and 90-140 Hz as fast gamma [45]. However, gamma oscillations are frequently nested in theta oscillations [46], whereas ripples are nested in sharp waves [47]. The frequencies observed in our carefully calibrated model of layer 2/3 mEC PV+ interneuronal networks consistently fall in the upper end of that range (∼150-200 Hz). This value is consistent with the frequency of gamma oscillations evoked in a study that selectively activated PV+ neurons at theta frequencies. It is possible that these high frequency oscillations are more analogous to the ripples in superficial mEC that contribute to ripple bursts and extended replays in area CA1 in quiet awake rodents [43]. Perhaps neural populations such as the stellate, pyramidal or other inhibitory neurons contribute to slowing the oscillations into a fast gamma range via synaptic or modulatory mechanisms. It has been previously suggested [48] that fast gamma oscillations are more similar to ripples than to slow gamma, and result from interneurons escaping control of phasic excitation and entering a regime of tonic excitation. In the former, the interneurons do not fire unless prompted by the phasic excitation, hence their intrinsic dynamics only contribute to setting the frequency in the tonic excitation regime. This interpretation is not universally accepted, however, as an alternative hypothesis posits that the ripples are merely transients resulting from a strongly synchronizing input [49].

### Relationship to Previous Models of Fast Oscillations in Heterogeneous Inhibitory Interneuronal Networks of Coupled Oscillators

Synchrony mediated by inhibition was originally thought to require a synaptic rise time longer than the duration of an action potential [50]. A pioneering study on the effects of heterogeneity in networks of interneurons generating fast oscillations implemented heterogeneity by simply changing the bias current of model neurons that were otherwise identical [11]. In the absence of conduction delays, synchrony was only observed for hyperpolarizing (but not shunting) synapses, and even then only for modest levels of heterogeneity in the bias current. A subsequent computational study [15] also implemented heterogeneity through the bias current; this study included conduction delays, and used stronger and faster synapses compared to the earlier study. Under these conditions, they found that shunting inhibition conferred greater robustness of synchronization at gamma frequency to heterogeneity in the excitatory drive than hyperpolarizing inhibition. Neurons with different levels of bias current traversed a different range of membrane potentials during the interspike interval (the more depolarized the bias current, the more depolarized the envelope). The effect of synaptic input with a shunting reversal potential was therefore different for different single interneurons. Specifically, it caused a phase advance in the slower, less depolarized neurons that increases the spike frequency, while leading to a phase delay when applied to the faster, more depolarized neurons that lowers the frequency. Therefore, shunting inhibition homogenized the rates, pulling them toward the center of the range. Although this study focused on interneuronal models with type 1 excitability, they reached similar conclusions using type 2 model neurons [51]. Our previous work on theta-nested gamma oscillations in inhibitory networks [52] also implemented heterogeneity using different levels of bias current; we found tighter phase locking of gamma oscillations to the theta modulation in type 2 models when using hyperpolarizing inhibition, and in type 1 models with shunting inhibition. Whether hyperpolarizing or shunting inhibition is more synchronizing and potentially more robust to heterogeneity will likely to depend on the exact model and synaptic parameters.

### Phase Response Curves

Phase response curve analysis provides a potential mechanism by which conduction delays or slow synaptic rise times, which can have a similar effect in stabilizing synchrony. Phase responses to strong inhibition often contain destabilizing discontinuities near a phase of 0 (or 1) [20,31,53,54], see Figure 5A (solid red and green traces). A conduction delay of sufficient duration can prevent noise from causing neurons to receive inputs on opposite sides of the discontinuity and stabilizes synchrony [26, 31]. In addition, in some cases, there may be an initial region of negative, destabilizing slope (Figure 5A, solid green trace at early phases). The region is destabilizing because if noise speeds up a neuron, accelerating its trajectory such that it receives an input from the population at a phase later than the 1:1 locking phase, the input further speeds the trajectory by advancing the time of the next spike. A slope of one is maximally stabilizing in this scenario because the phase response exactly compensates for the input arriving later (earlier) by delaying (advancing) the spike by exactly the difference between when the input actually arrived, and when it would have arrived in a synchronous mode [27,28,55]. The slope of the phase resetting curve at the locking point can also provide insight into the speed at which the network will synchronize; if it is flat, synchrony can only be weakly attracting and easily disrupted by noise. Another insight is that a steeper PRC has a larger range of advances and delays available so that it is more likely to be able to adjust its own frequency to match that of the population.

### Caveats on generality

Our study is specific to layer 2/3 medial entorhinal cortex because both intrinsic and synaptic parameters of the network were constrained by data from this region [32]. The only other study that has previously attempted to capture the true extent of heterogeneity in the intrinsic properties of the PV+ interneuronal network, in the external segment of the globus pallidus [58], did not address network synchronization. Although we have strong evidence for type 2 excitability of PV+ neurons in the MEC [20, 23], previous work supports type 1 excitability in the globus pallidus [59], substantia nigra pars reticulate [60], dentate gyrus [61] and hippocampal area CA1 [62–64]. Moreover, the PV+ neurons in both the globus pallidus and the substantia nigra pars reticulata are spontaneous pacemakers in a slice preparation [65], whereas PV+ neurons in the other regions are quiescent. The synchronization properties of different brain circuits containing inhibitory PV+ networks are not generic and likely differ greatly between brain regions.

### Coupled oscillators versus stochastic population oscillators

Two modes of neural synchrony have been proposed [44,66,67]: a strong synchrony in which coupled oscillators fire on every cycle, and a weak synchrony that arises from the population dynamics in which the firing of individual neurons is sparse and appears stochastic. For strong synchrony, most if not all of the recruited neurons fire on almost every cycle of the network oscillation, and the interspike interval histogram has a sharp peak at the network frequency, possibly with subharmonic peaks indicating skipped cycles. For weak synchrony, neurons fire sparsely and irregularly with only a few neurons participating in any given cycle of the network oscillation. Despite clear peaks in the spike density at the population period, the sparseness can obscure any peaks in the ISI histogram of individual neurons such that it resembles a left-truncated exponential distribution characteristic of a Poisson process with a refractory period.

We are not aware of single unit recordings in the mEC during ripples, but in area CA1, PV+ neurons fire at 122±32 Hz [68] during ripples *in vivo,* with PV+ basket cells discharging on virtually every ripple event [36,69,70]. Such high firing rates are clearly not consistent [8] with a stochastic population oscillator [67], and our model in this study of theta-nested fast oscillations is clearly a coupled oscillator model. A recent computational study [71] on ripple generation in area CA1 found that, in some cases, an inhibitory interneuronal population can exhibit strong synchrony, while the excitatory neuron population simultaneously exhibits weak stochastic synchrony. That study modeled single neurons as conductance-based leaky integrate-and-fire neurons and assumed that fast oscillations are a network dynamical pattern that does not crucially depend on the details of subthreshold dynamics and spike generation. In contrast, we hypothesize that the details of subthreshold dynamics and spike generation crucially affect synchronization via their phase response tendencies, which can differ greatly from those of leaky integrate and fire neurons [44].

## Methods

### Ethics statement

All experimental protocols were approved by the Boston University Institutional Animal Care and Use Committee.

### Experimental Methods

Methods were as in [32], with some data taken from recordings collected in that study; here is a brief summary. Resting membrane potential was obtained by averaging across 1 s of the recorded membrane potential in the absence of an external input. Input resistance was calculated using the inverse of the slope of the linear fit to the steady state current voltage (I-V) relationship measured between 0 and 100 pA of injected current in 25 pA increments. The membrane time constant was obtained by fitting the voltage trace to a single exponential during the return to RMP after a 100 pA hyperpolarizing current step. For frequency-current measures, a series of depolarizing current steps of 25 pA were used to depolarize neurons and drive action potential generation. Some of the recorded neurons generated a weak early spike frequency adaptation, which was replicated in the model by accumulation of Na inactivation and Kv1 activation within a few successive spikes (Fig. 2B). The f/I curves shown in Figure 2C show the steady-state frequency after a weak early spike frequency adaptation. Only values at which repetitive firing could be sustained were plotted. Recorded neurons also presented a much weaker and slower later adaptation, likely due to A-type potassium currents [79], which we omitted from the model for simplicity. For gap junction measures, a square hyperpolarizing pulse between -100 and -300 pA was used to hyperpolarize the pre-synaptic cell across 25-50 trials that were averaged in the post-synaptic cell. A measured junction potential of ∼11 mV was not subtracted from recordings. Recordings were taken from slices between 3.2 mm and 4.3 mm from the dorsal surface (bregma) of the brain. C57BL/6J background, PV-Cre mice [72] (Jackson Labs, stock # 017320) were crossed with the lox-stop-lox tdTomato reporter mice [73] (Jackson Labs, stock # 007914) to visualize PV+ interneurons. Eleven representative dorsal mEC PV+ neurons were utilized for the calibrations shown in Figures 1 and 2.

### Computational Methods

Although a single PV+ basket cell in region CA1 makes chemical synaptic contacts with about 60 other PV+ basket cells [74], we are unaware of similar data in the mEC. For this reason this number is not well-constrained in our model. In the mEC, the probability of both electrical and chemical synapses drops off dramatically at distances between somata that are 125-150 μm apart [32]. In order to apply the connection probabilities, the average number of PV+ cells that are within that distance of a typical soma of a PV+ cell must be estimated. This estimate can be refined by the size of the area illuminated by the laser. The diameter of this area is about 200 μm [12]. Thus, we conservatively estimated that each cell makes 36 chemical synaptic contacts onto other PV cells activated by the ChR2, with a maximum of 27 electrical contacts, which aligns with the measured connectivity probabilities if we assume a network of 100 neurons.

All simulations were carried out in the BRIAN simulator [75]. The simulation code has been uploaded to modelDB at https://senselab.med.yale.edu/modeldb/enter-Code?model=267338#tabs-1 with password PVfs1n3t . The network consists of 100 single compartment model neurons with five state variables: the membrane potential, *V*, and four gating variables (m, h, n, and a) that use the same kinetic equations as the original Hodgkin-Huxley model [76, 77], but with different parameters tuned to replicate the dynamics of fast spiking neurons in the mEC. Also, consistent with other models of fastspiking interneurons [51, 78], we included two delayed rectifier K^+^ currents (I_Kv1_ and I_Kv3_) . The differential equation for the membrane potential (V) of each neuron is *C_M_ dV* / *dt* = *I_app_* − *I_Na_* − *I_Kv_*_1_ − *I_Kv_*_3_ − *I_L_* − *I_syn_* − *I_gap_* − *I_ChR_*, where C_M_ is the membrane capacitance, I_app_ is an externally applied current that is only nonzero when simulating step currents for electrophysiological measurements, I_Na_ is the fast sodium current, I_L_ is the passive leak current, I_syn_ is the GABA_A_ synaptic current, I_gap_ is the gap junctional current and I_ChR_ is the simulated sinusoidal optogenetic drive. The equations for the intrinsic ionic currents are as follows: *I_Na_*= *g_Na_ m*^3^*h*(*E_Na_*−*V*), *I_Kv_*_1_ = *g_Kv_*_1_ *a*^4^ (*E_K_*−*V*), *I* = *g_Kv_*_3_ *n*^4^(*E_K_*−*V*) and *I_L_*= *g_L_* (*E_K_* −*V*), with E_Na_= 50 mV, E_K_= -90 mV and E_L_ varied across the population.

The dynamics of the gating variables are given by *dx* / *dt* = *α_x_* (1− *x*) − *β_x_ x* for the activation variables (m, n, a) and by *dx* / *dt* = *β_x_* (1− *x*) − *α_x_ x* for the inactivation variable h, where *α_x_* = *k*_1*x*_ (*θ_x_* −*V*) / (exp((*θ_x_* −*V*) / *σ*_1*x*_) −1) and *β_x_* = *k*_2 *x*_ exp(*V* / *σ* _2 *x*_) using parameters in Table1.

The simulated f/I curves in Figure 2C2-C3 and the simulations without gap junctions in Figures 4, 6A1 and 6B1 did not include gap junctions. In simulations without gap junctions, the leakage reversal potential E_L_ was set to the resting membrane potential. Values for the passive parameters in these neurons were generated from distributions with ranges similar to the experimental ones, as shown in Figure 1C.

Some of the parameters (see Table 1) for the voltage-gated currents were uniform across the 100 simulated neurons. The remaining parameters for the active properties were also generated from uniform random distributions (range): g_Na_ (6000,35000) nS, g_Kv1_ (15,150) nS, θ_m_ (-60,-45) mV, θ_h_ (-60, -50) mV, θ_n_ (-15, 25) mV, θ_a_ (35, 55) mV. The Kv3 peak conductance was set randomly to between 3 and 5% of the peak Na conductance, which helped reproduce some of the features from the experimentally recorded traces. These features included the AP waveform (including peak height, AHP depth, and peak width), transient spikes observed near the onset of step current for near-rheobase amplitudes (see example in Figure 2AB), and f/I curve ranges (including rheobases and cutoff frequencies), as shown in Figure 2C. Parameter sets that did not reproduce the desired features were discarded, and additional sets were generated until 100 unique parameters sets were obtained for each of the 100 model neurons.

The connection probabilities and conductances for both electrical and chemical synapses were taken from [32]. Chemical synapses were modeled by an inhibitory postsynaptic conductance with a biexponential waveform *g_i,k_* (*t*) = *F* (exp(−(*t* − *t_i_^k^*− *δ_i_*) / *τ*_1_) −exp(−(*t* − *t_i_^k^* − *δ_i_*) / *τ*_2_)) where F is a normalization factor that sets the peak to one [80]. This conductance waveform was initiated after a random, uniformly distributed delay, δ, ranging from 0.6 to 1.0 ms by each spike *k* in the presynaptic neuron *i*:*I_syn_* = ∑*_i_* ∑*_k_ g_syn_*_,*i*_*g_i,k_* (*t*)(*E_syn_* −*V*) . The delays, as well as a fixed τ_1_ of 0.484 ms (which corresponds to a rise time of 0.4 ms, see [80]) and a fixed decay time constant τ_1_ of 2.3 ms, were calibrated according to the experimental data in [19]. The reversal potential E_syn_ was varied to simulate shunting (-55 mV) and hyperpolarizing inhibition (- 75 mV). The maximal synaptic conductance was lognormally distributed [57] with parameters μ=0 corresponding to the log of 1 nS and σ = 1. The probability of connection in each direction was 0.36.

The gap junction current is given by *I_gap_* = _∑*i*_ *g_gap_*_,*i*_ (*V* −*V_i_*) summed over the *i* neurons connected to a given neuron. When gap junctions are included, the measured input resistance is no longer determined by the leakage conductance alone, but also by the gap junctional conductance. To first order, one can approximate the input resistance for each neuron *i* by *R_input_*_,*i*_ =1/ (*g_L_* + ∑*_i_ g_gap_*_,*i*_) . We used a 27% probability that any pair of neurons was connected by gap junctions [32]. Peak conductances were obtained from data originally collected for [32]. The histogram of peak conductances for the electrical synapses in Figure 3B suggests a bi-modal distribution with a weak mode given by the positive half of a Gaussian distribution with zero mean and a standard deviation of 0.4 nS, and a strong mode, which we approximated by a Dirac delta at 1.2 nS. For each neuron *i* in the network, we chose 27 other neurons at random, labeled *j*. We then assigned a 25% probability that a particular connection was of the strong mode type, and 75% that it was of the weak mode type. For strong mode synapses, a value of 1.2 nS was assigned tos *g_ji_^gap^* whereas for the weak mode it was drawn from from the truncated Gaussian with st.d. 0.4 nS. Each gap junction was added bidirectionally with the same strength, and *g_ji_^gap^* was subtracted from the leakage conductance of both neurons to keep R_input_ in the experimentally constrained range. Gap junctions were only added while the leakage conductance remained above a predefined floor value g_L_^min^ (1.5 nS), since this quantity cannot be zero or negative. Frequently the total number of electrical synapses was less than 27 for a given neuron, but this procedure was necessary to honor the data since passive properties were measured with gap junctions intact. The network synchrony and dominant frequency were not very sensitive to this parameter for values between 1 and 2 nS (Figure S4). The capacitance was not changed because the input resistance was approximately preserved. Leakage reversal potentials were then adjusted to preserve the experimentally constrained distribution of resting membrane potentials.

The optogenetic drive is present in the network simulations as *I* = *g_ChR_* sin(2*π ft* /1000)(*E_ChR_* −*V*), where t is in ms, f is 8 Hz, E_ChR_ is 0 mV, and g_ChR_ is 14 nS.

Synaptic depression was calibrated according to [32] using the model by Markram and Tsodyks [81] adapted by [82], but neglecting facilitation. In this model, the available fraction of transmitter x evolves according to: 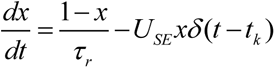, where t_k_ is the *k*th spike time, τ_r_ =100 ms is the recovery time to replenish the available pool of vesicles for release, U_SE_ =0.3 is the fraction of available pool released by each spike, and the value of x prior to a spike is proportional to the peak current value of the inhibitory post-synaptic current.

A forward Euler method was used to integrate the ODEs. For the calibration of passive and active properties, and for the network simulations without gap junctions, an integration time step of 0.01 ms was used. Simulations did not converge with that time step when heterogeneous gap junction conductances were considered, and the time step was reduced to 5.0e-4 ms. The histograms in Figures 4, 6 and 8 were computed using a time step of 0.1 ms.

The frequency, power and theta phase onset and offset for the fast oscillations were obtained from Complex Wavelet Transform powers like the example shown in Figure8C1. The wavelet transforms were computed using the *cwt* function from *scipy.signal* with a Morlet wavelet (*Morlet2* in *scipy.signal*) of order 5 applied over a population rate. In particular, the wavelet scalings were set as *order*\*sampling_rate/(2*π\**f*). The wavelet power was obtained as the squared modulus of the complex wavelet transform. The population rate used to compute it was obtained using a flat sliding window of width 0.1 ms (*sampling_rate*) in the function *PopulationRateMonitor.smooth_rate()* from BRIAN 2. The considered range of frequencies were from *f*=50 to 449 Hz in steps of 3 Hz. No normalization was applied.

The dominant frequency, f_max_, or frequency of maximum power in Figure 7, was computed from all theta cycles for each connectivity pattern as the frequency with maximum wavelet. The mean and standard deviation of the dominant frequency were computed only from cycles whose power was above 0.3 times the maximum power recorded for that connectivity pattern in order to avoid contamination by cycles in which no synchrony was observed.

The circular histograms for onset and offset theta phases of the fast oscillations were computed using the *rose.diag* function in the *circular* package from CRAN R by pooling together the 900 values corresponding to each parameter set, i.e. the 30 last simulated theta cycles for each of the 30 networks with different connectivity pattern. The first four cycles were discarded to eliminate transients.

---

**Figure S1.**
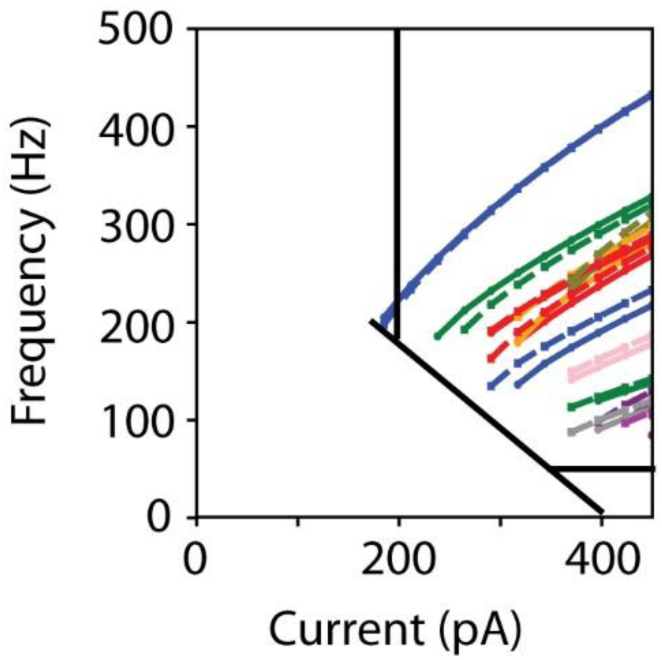
f/I curves for model network with gap junctions intact. Curves for fifteen representative model neurons are shown.

**Figure S2.**
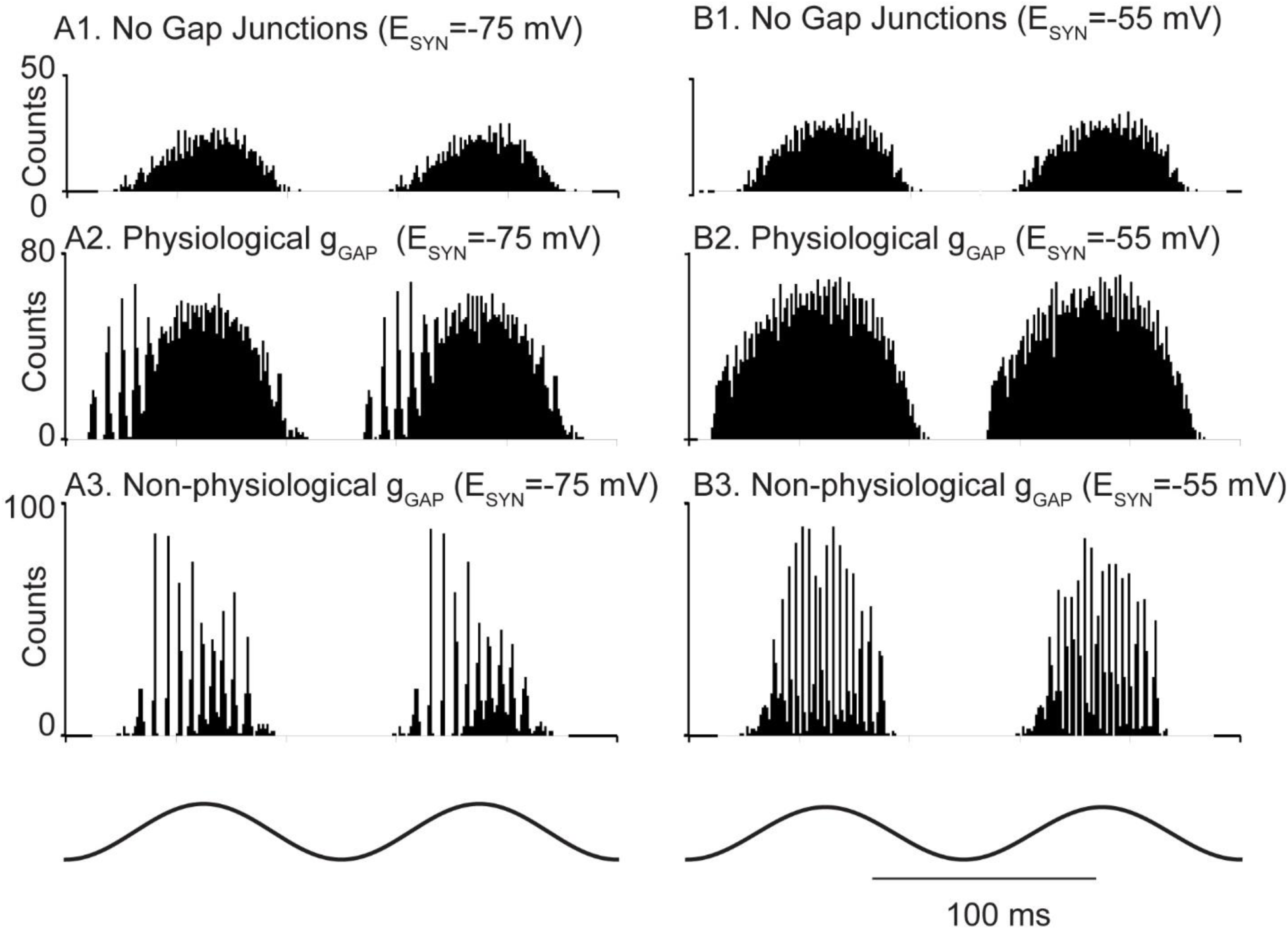
The simulations in Figure 6 were rerun with conduction delays set uniformly to 1.6 ms with very similar results.

**Figure S3.**
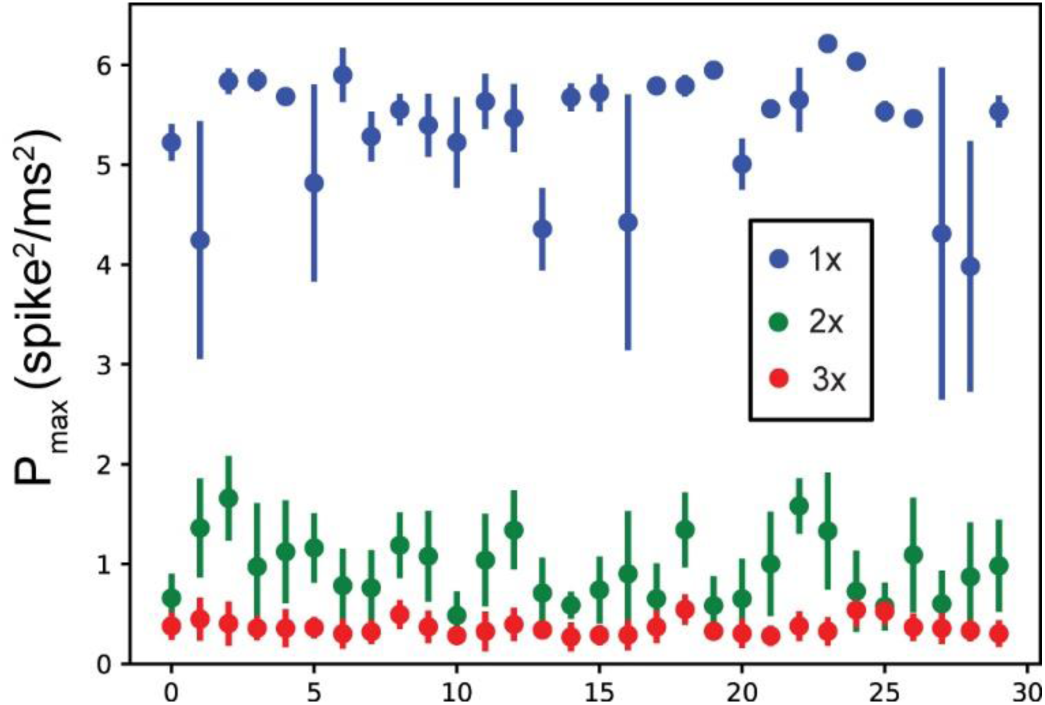
Effect of increasing synapse strength in shunting networks with unphysiologically strong gap junctions.

**Figure S4.**
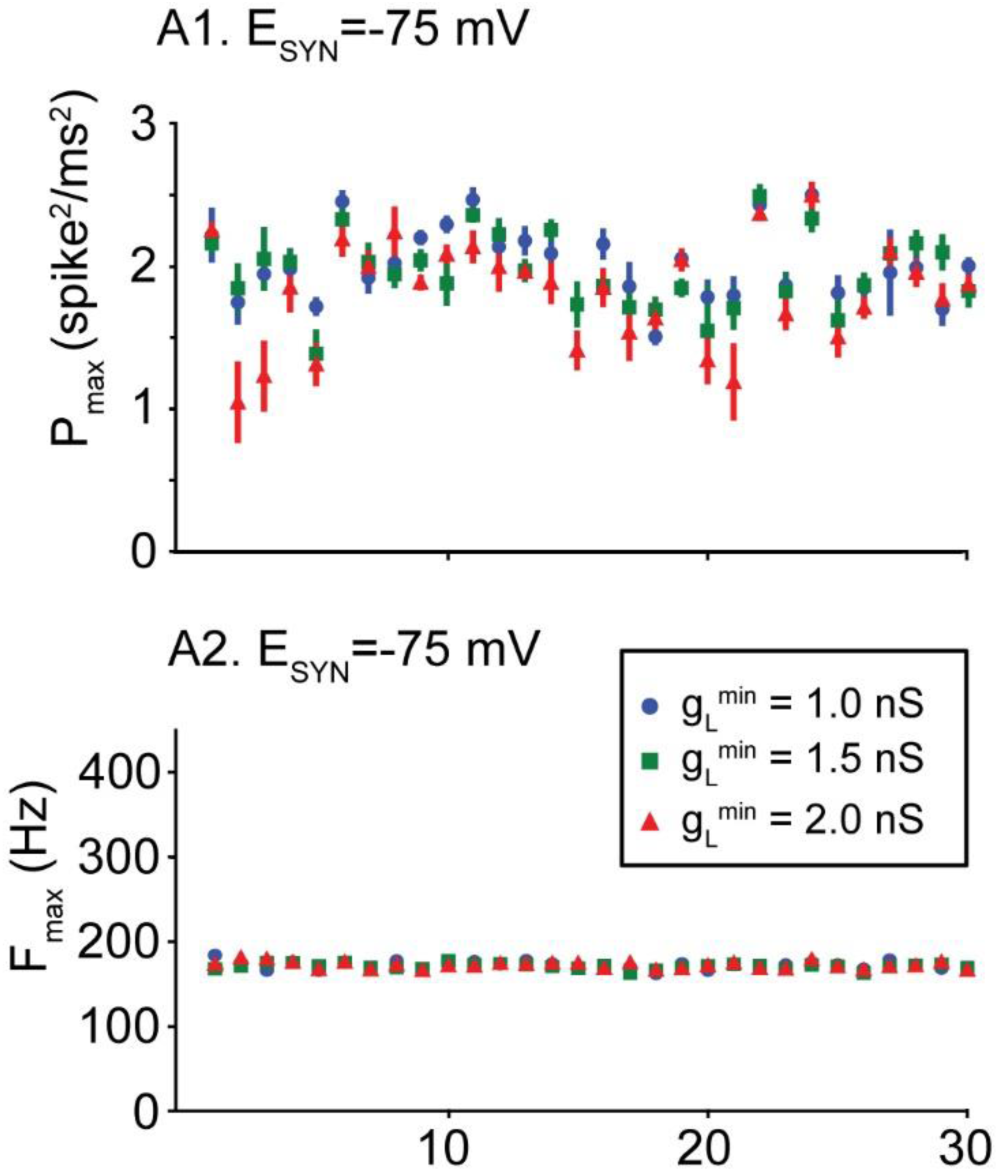
Values for g_L_ floor do not substantially affect oscillatory frequency or power.

